# Common anesthetic used in preclinical PET imaging inhibits metabolism of the PET tracer [^18^F]3F4AP

**DOI:** 10.1101/2023.12.14.571690

**Authors:** Karla Ramos-Torres, Yang Sun, Kazue Takahashi, Yu-Peng Zhou, Pedro Brugarolas

**Author notes:** **Correspondence:** Pedro Brugarolas, PhD, 55 Fruit St, Bulfinch 051, Boston, MA 02114.

## Abstract

PET imaging studies in laboratory animals are almost always performed under isoflurane anesthesia to ensure that the subject stays still during the image acquisition. Isoflurane is effective, safe, and easy to use, and it is generally assumed to not have an impact on the imaging results. Motivated by marked differences observed in [^18^F]3F4AP brain uptake and metabolism between human and nonhuman primate studies, this study investigates the possible effect of isoflurane on [^18^F]3F4AP metabolism and brain uptake. Isoflurane was found to largely abolish tracer metabolism in mice resulting in a 3.3-fold higher brain uptake in anesthetized mice at 35 min post radiotracer administration, which replicated the observed effect in unanesthetized humans and anesthetized monkeys. This effect is attributed to isoflurane’s interference in the CYP2E1-mediated breakdown of [^18^F]3F4AP, which was confirmed by reproducing a higher brain uptake and metabolic stability upon treatment with the known CYP2E1 inhibitor disulfiram. These findings underscore the critical need to examine the effect of isoflurane in PET imaging studies before translating tracers to humans that will be scanned without anesthesia.

## INTRODUCTION

Positron emission tomography (PET) is a molecular imaging technique that uses biologically active radiolabeled compounds to visualize and quantify biochemical processes inside the body^1^. PET is widely used in oncology^2^, neurology^3^, cardiology^4^, and drug discovery^5^. Furthermore, in recent decades there has been remarkable growth in the development of novel tracers for emerging targets^6,7^. By detecting high-energy photons produced in the positron-electron annihilation, PET can provide quantitative images of a radiotracer distribution deep inside the body with high biochemical specificity and sensitivity. Given these properties, new tracers can be tested in animal models with reasonable certainty that the results in will translate to humans with good reproducibility^8^.

3-[^18^F]fluoro-4-aminopyridine ([^18^F]3F4AP) is a radiolabeled derivative of the FDA-approved multiple sclerosis drug 4-aminopyridine (4AP, dalfampridine) developed for imaging demyelinated axons in the central nervous system by PET^9^ (**Fig. 1**). Similar to 4AP, [^18^F]3F4AP binds to voltage-gated K^+^ channels in the brain, including exposed K_v_ channels in demyelinated axons. Previous work has shown that [^18^F]3F4AP is very sensitive to demyelinated lesions in rodents and nonhuman primates (NHPs)^9,10^. Preclinical evaluation in NHPs presented fast brain uptake (SUV > 3 at 4 min) and washout, high plasma availability, and high metabolic stability (>90% parent fraction up to 2h post injection)^10^. Due to its potential for detecting demyelination this tracer has recently advanced into human studies (clinicaltrials.gov: NCT04699747, NCT04710550). First-in-human imaging with [^18^F]3F4AP revealed widespread biodistribution and high brain penetration, but also exhibited faster than expected clearance rate and greater metabolism of the radiotracer in blood (less than 50% parent remaining 1h post injection)^11^. Given that the metabolic breakdown of the tracer may lead to a reduced brain uptake, introduce background signal, and create variability across subjects, further studies are warranted. Moreover, the unforeseen difference in metabolic stability between humans and NHPs, which is normally similar^12^, is intriguing.

**Figure 1.**
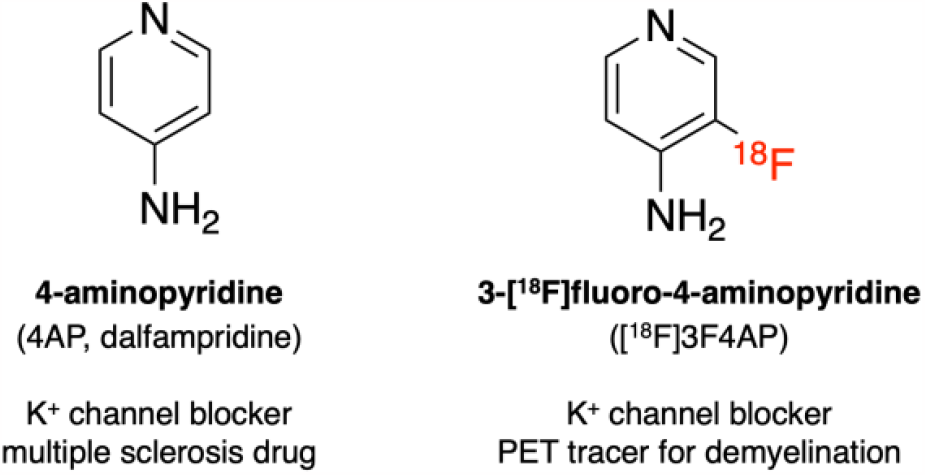
Chemical structures of 4AP and [^18^F]3F4AP.

The pharmacokinetics of 4AP have been carefully evaluated in a variety of species including rats^13^, dogs^14^, horses^15^, cattle^16^, and humans^17^. Studies following oral administration of 4AP in rats, dogs and humans have shown rapid absorption and renal clearance of the drug primarily as an unchanged compound^13,18,19^. Characterization across species has reported limited metabolism of 4AP. In humans 24h after oral administration of ^14^C-labeled 4AP ∼90% of the radioactive species recovered in urine corresponded to the unchanged drug, with the remaining corresponding to the 3-hydroxy derivative and its sulfate conjugate^20^. Investigation of the metabolic pathways of 4AP points to an initial step catalyzed by members of the cytochrome P450 family, where CYP2E1 has been identified as the key metabolic enzyme.

CYP2E1 is the most abundant isoform of cytochrome P450 in the human liver^21^. P450 enzymes are responsible for the oxidation of over 90% of drugs^22^. CYP2E1 plays an integral part in the breakdown and clearance of a variety of small molecules including acetaminophen and halogenated anesthetics^23^. Recent *in vitro* studies from our laboratory have shown that CYP2E1 is also able to metabolize (nonradioactive) 3F4AP very efficiently^24^. In an *in vitro* competition assay the affinity of CYP2E1 for 3F4AP was 50 times higher than the affinity for 4AP, suggesting a primary role for CYP2E1 in metabolizing 3F4AP. Moreover, CYP2E1 is the predominant cytochrome P450 isoform responsible for human clinical isoflurane metabolism *in vivo*^25^. Given these observations, we postulated that isoflurane anesthesia could play an interfering role in the observed differences in uptake, clearance, and metabolic breakdown of [^18^F]3F4AP between monkeys and humans. The goals of this study were to (i) determine whether isoflurane affects the *in vivo* stability of [^18^F]3F4AP; (ii) examine the formation and presence of radioactive metabolites of [^18^F]3F4AP in the brain; and (iii) inhibit the metabolism of [^18^F]3F4AP *in vivo* by introducing clinically approved CYP2E1 inhibitors, such as disulfiram.

## METHODS

### Compliance

All rodent procedures were approved by the Institutional Animal Care and Use Committee (IACUC) at the Massachusetts General Hospital. All animal studies were conducted in compliance with the ARRIVE guidelines (Animal Research: Reporting in Vivo Experiments) for reporting animal experiments.

### Radiochemistry

[^18^F]3F4AP was produced in a GE TRACERlab Fx2N synthesizer according to previously reported procedure^26^. Semipreparative HPLC separations were performed on Sykam S1122 Solvent Delivery System HPLC pump with the UV detector at 254 nm with a Waters C18 preparative column (XBridge BEH C18 OBD Prep Column 130 Å, 5 µm, 10 mm × 250 mm). The obtained [^18^F]3F4AP fraction (radiochemical purity > 99%, n = 5), in 95% 20 mM sodium phosphate buffer, pH 8.0, 5% EtOH solution) passed the QC (with Thermo Scientific Dionex Ultimate 3000 UHPLC equipped with Waters XBridge BEH C18 analytical column [130 Å, 3.5 µm, 4.6 × 100 mm], 95% 10 mM sodium phosphate buffer, pH 8.0, 5% EtOH solution as mobile phase, t_R_ = 4.7 min) and was ready for injection into animals.

### Evaluation of anesthesia effects on the radioactivity concentration in whole blood and brain

Two cohorts of mice (9-12 week-old, male) were administered 50-100 µCi of [^18^F]3F4AP intravenously via tail vein injection. Radiotracer delivery was performed under isoflurane anesthesia (1.5 – 2 % isoflurane in O_2_) for the first cohort (n = 8), which was kept anesthetized for the duration of the experiment. For the second cohort (n = 9), mice were temporarily physically restrained while awake for radiotracer administration and then allowed to move freely. For both groups, 35 minutes post-injection of the radioactive dose mice were euthanized by intraperitoneal injection of pentobarbital sodium and phenytoin sodium solution (Euthasol^®^). Brain tissue was harvested, and blood was collected via cardiac puncture to measure radioactivity concentration via gamma counting. Gamma counting was performed in a PerkinElmer 2480 Wizard Automatic Gamma Counter and calibration curve was produced with [^18^F]3F4AP standards to calculate radioactivity concentration in biological samples.

### Evaluation of anesthesia effects on the *in vivo* metabolism of [^18^F]3F4AP

Two cohorts of mice (9-12 week-old, male) were administered 250-300 µCi of [^18^F]3F4AP intravenously under isoflurane anesthesia (n = 11) or temporary physical restriction while awake (n =11). For both groups, 35 minutes post-injection of the radioactive dose mice were euthanized by Euthasol^®^ intraperitoneal injection. Blood was collected via cardiac puncture to measure radioactivity concentration in whole blood (WB) and plasma (PL) via gamma counting. Brain tissue was harvested, mixed with 1.0 mL (1:2 w/v) freshly prepared 0.4 N perchloric acid (HClO_4_) aqueous solution, and mechanically homogenized via ultrasonication at 30,000 rpm for 30 seconds at 4 ºC with a Dremel^®^ Moto-Tool. Homogenized lysates were centrifuged at 21000 RCF (8 ºC) for 7 min. Resulting supernatant was collected for polar radiometabolite analysis. Radioactivity concentrations of whole brain, homogenized lysates, supernatant, and residual pellet were measured via gamma counting. All radioactivity concentrations were measured using a single well high-purity germanium gamma detector. All blood, plasma and tissue samples were kept on ice at 0 ºC for the duration of the experiment.

### Radio-HPLC analysis of metabolites in plasma and brain samples

Plasma samples and pH-neutralized brain supernatant samples were first filtrated through a 10K filter (Amicon™ Ultracel-10 regenerated cellulose membrane, 0.5 mL sample volume, Centrifugal Filter Unit) by centrifugation at 21000 RCF (8 ºC) for 15 min. The filtrates were injected into HPLC (Agilent 1260 Infinity II, along with Eckert & Ziegler FlowCount radioisotope detector) through a C18 column (XBridge, BEH, 130 Å, 3.5 µm, 4.6 × 100 mm) with a vanguard cartridge (XBridge, BEH, 130 Å, 3.5 µm, 3.9 × 5 mm) using 10 mM NH_4_HCO_3_ aqueous solution (pH 8.0) and MeCN (96:4) as mobile phase.

### *In vivo* inhibition of [^18^F]3F4AP metabolism by disulfiram treatment

### Preparation and administration of disulfiram sesame oil suspension

Disulfiram (Sigma Aldrich) sesame oil suspension was prepared freshly as a 6 mg/mL stock solution. Each mouse was administered 40 mg/kg disulfiram via intraperitoneal injection 2 hours prior to the radioactive dose administration.

### Analysis of the effects of disulfiram treatment on the *in vivo* uptake and metabolism of [^18^F]3F4AP

Following pre-treatment of two cohorts of mice (10-week-old, male) with disulfiram (n = 6) or sesame oil vehicle control (n = 4), 250-300 µCi of [^18^F]3F4AP were administered intravenously under temporary physical restriction while awake. Mice were euthanized 35 min post-tracer administration, blood and brain was collected, processed, and evaluated *ex vivo* for radioactivity content by gamma counting and metabolite content by radio-HPLC.

### Statistics

Descriptive statistics including mean, standard deviation (SD) and standard error (SE) were calculated for each group. Two-group t-tests and multi-group t-tests (e.g., ANOVA) with a significance level of α = 0.05 were used to assess differences among groups. Grouped data are reported as mean ± SE.

## RESULTS

### Study design

To test our hypothesis that isoflurane anesthesia interferes with the *in vivo* breakdown of [^18^F]3F4AP, we wanted to investigate metabolism of the tracer in awake and anesthesia conditions within the same species. Given the challenges and ethical considerations to test awake rhesus macaques or anesthetized humans, we opted to do the investigation in awake and anesthetized mice. Awake mice received the radiotracer dose while momentarily restrained and were returned to their home cage until the time of euthanasia whereas anesthetized mice received the dose under isoflurane and were kept anesthetized for the duration of the experiment (**Fig. 2**). Euthanasia at 35 min post tracer administration was empirically chosen as an approximation to an equivalent time point of 90 min post injection in NHPs and humans. At this time, the initial phase of brain time-activity curves (TACs), characterized by a fast increase and rapid washout, has passed and brain radioactivity has reached the plateau/terminal stage^9,10^. After euthanasia, blood was collected via cardiac puncture and the brain dissected to measure radioactivity concentration and analyze radiometabolites present in these tissues. Although not perfusing the animals meant that blood would be present in the brain sample (around 5% of brain volume corresponds to blood) we decided to not perfuse to avoid washing out some of the loosely bound metabolites within the brain. To measure the metabolites in blood, the plasma was separated from the cellular components using standard centrifugation techniques and the proteins and other large macromolecules filtered out using 10 kDa centrifugal filter prior to injection onto the radioHPLC. From our experience with [^18^F]3F4AP blood radiometabolite analysis, this method is superior to trapping the radioactive species in a solid phase extraction cartridge followed by elution and injection onto the HPLC. To measure the radiometabolites in brain, the brain was mechanically homogenized in perchloric acid and centrifuged at high speed to separate solid components from soluble components. In our tests, this method was superior to precipitating and extracting the organic components with acetonitrile and resulted in extraction of 69 ± 2% of radioactivity in the supernatant. The supernatant was then filtered using the same 10 kDa centrifugal filter and the pass-through neutralized with base prior to injection on the HPLC. These methods had the added advantage that no organic solvent was introduced which would compromise the retention of the tracer and any polar radiometabolites on the HPLC column.

**Figure 2.**
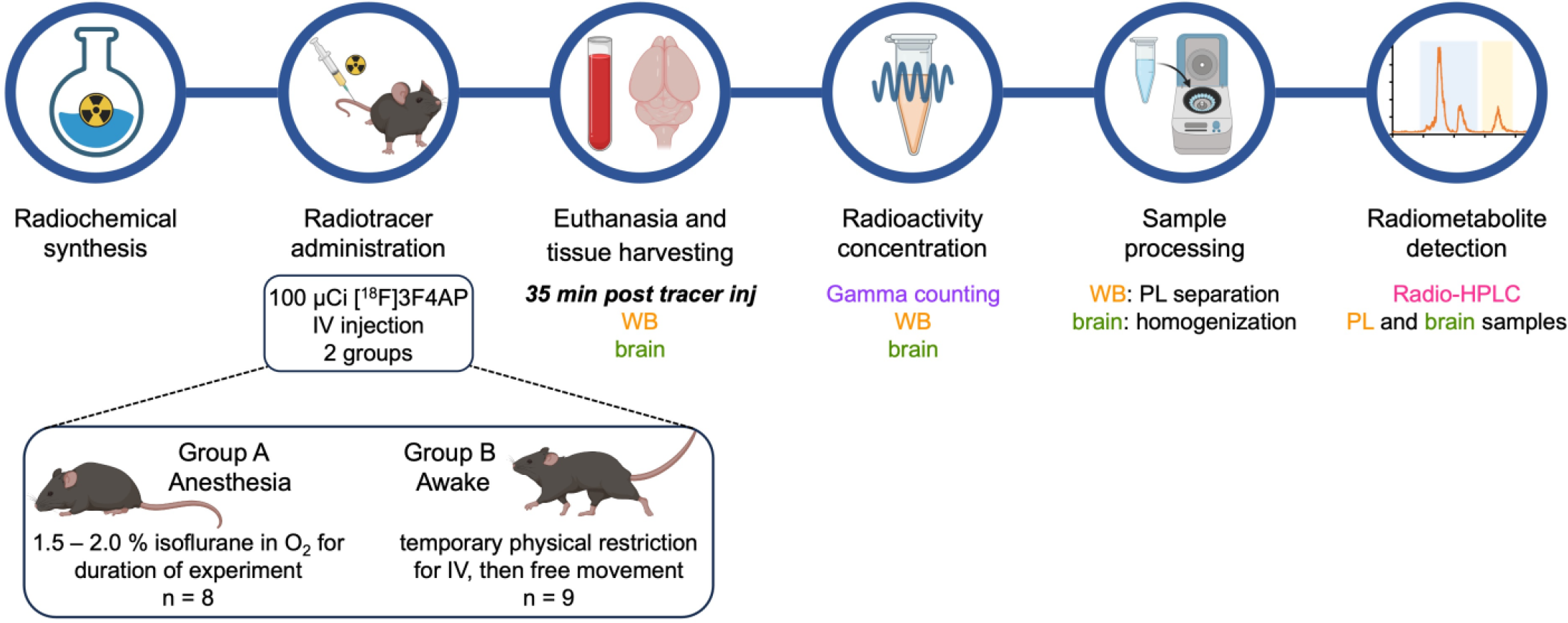
Experimental design. Young adult mice are assigned to two groups (isoflurane and awake) at random. 35 min after intravenous administration of [^18^F]3F4AP the animals are euthanized, their blood (WB) and brain collected, and radioactivity concentration assessed. Tissue samples are processed, and plasma (PL) and brain homogenates analyzed for radiometabolite content. Created with BioRender.com

### Isoflurane anesthesia results in higher [^18^F]3F4AP brain uptake

Recent studies have shown that [^18^F]3F4AP uptake is markedly different in anesthetized monkeys^10^ and awake human subjects^11^. Analysis of reported PET imaging results shows that at 90 minutes post-tracer administration, [^18^F]3F4AP uptake in the brain of anesthetized NHPs corresponds to an SUV_90 min-br(NHP)_ of 1.50 while human imaging studies in awake subjects show an SUV_90 min-br(human)_ of 0.54. Blood sampling performed during these studies show comparable radioactivity concentration in whole blood (WB) for both anesthetized NHPs and awake humans (SUV_90 min-WB(NHP)_ 0.946 *vs*. SUV_90 min-WB(human)_ 0.87) (**Fig. 3A**). To investigate whether isoflurane anesthesia could be playing a role in the observed difference in brain uptake and blood concentration of [^18^F]3F4AP, we compared radioactivity concentration in WB and brain of mice that had been administered the radioactive tracer while anesthetized or awake. As observed in **Figure 3B** mice that had received the dose under isoflurane anesthesia and were kept anesthetized showed higher radioactivity concentration in brain when compared to awake mice. At 35 min post-injection [^18^F]3F4AP brain uptake was 3.3-fold lower in awake mice in comparison to anesthetized mice (SUV_35 min-br(iso mice)_ 1.40 ± 0.13, n = 8 *vs*. SUV_35 min-br(awake mice)_ 0.42 ± 0.03, n = 9; *p* < .001). In contrast, no significant differences were found in WB SUV values between the two groups (SUV_35 min-WB(iso mice)_ 0.82 ± 0.07 *vs*. SUV_35 min-WB(awake mice)_ 0.77 ± 0.03). Interestingly, the observed brain and blood concentration in anesthetized and awake mice at 35 min post injection is remarkably similar to that of anesthetized NHPs and awake humans at 90 min.

**Figure 3.**
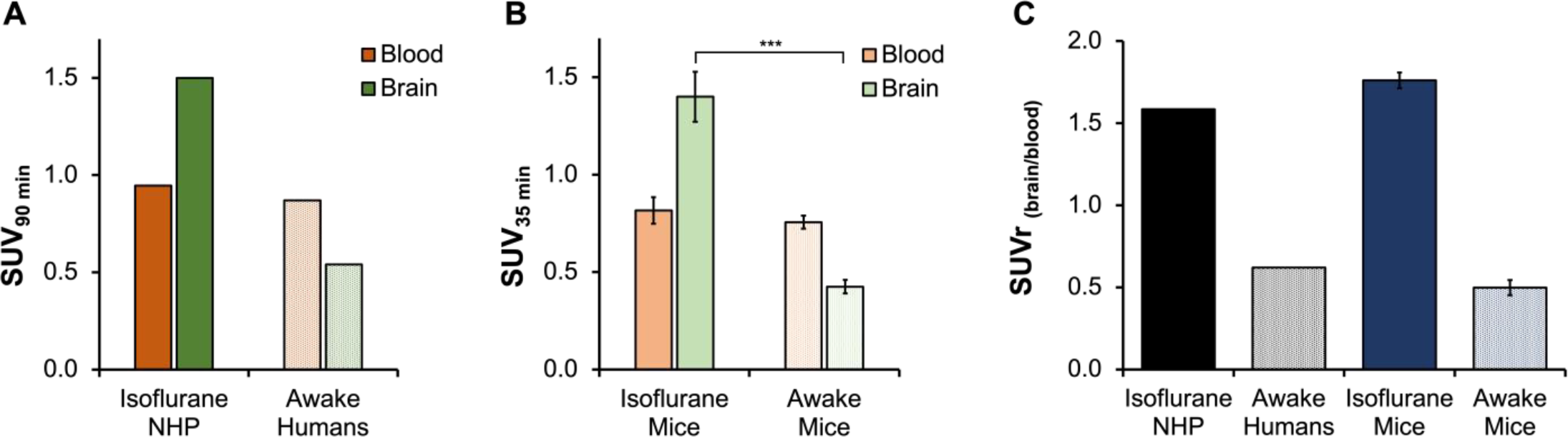
[^18^F]3F4AP uptake in whole blood and brain of anesthetized and awake subjects. A) Tracer uptake in anesthetized non-human primates (rhesus macaque) and awake humans in whole blood and brain at 90 min post-tracer administration. Values were calculated from previous reported experiments^10,11^. B) Tracer uptake in whole blood and brain samples of anesthetized (n = 8) and awake (n = 9) mice at 35 min post-tracer administration. C) Normalization by brain-blood ratios in anesthetized and awake subjects calculated from data in panels A and B.

### Isoflurane anesthesia reduces [^18^F]3F4AP metabolism

Upon confirmation that isoflurane anesthesia plays a role in the *in vivo* brain uptake of [^18^F]3F4AP, we set to evaluate the presence of radiometabolites in the plasma and brain of mice with and without isoflurane. We posited that the lower concentration in brain may be driven by metabolism of the tracer. As hypothesized, higher metabolism was observed in the plasma and brain of awake mice in comparison to anesthetized, revealed by the percent of parent fraction (%PF) remaining at 35 min post tracer administration. **Figure 4A** shows representative radio-HPLC traces of plasma and brain samples for each anesthesia condition. In plasma, anesthetized animals showed 74.8 ± 1.6 % of unmetabolized [^18^F]3F4AP remaining in circulation 35 minutes after intravenous administration, while awake animals presented only a 17.7 ± 1.7 % of the parent [^18^F]3F4AP. While polar radiometabolites accounted for a significant share of the radioactivity concentration in the plasma of both anesthetized and awake animals, in brain the parent tracer had the higher radioactivity contribution. For anesthetized mice, unmetabolized [^18^F]3F4AP was present in 98.4 ± 0.2 % in the brain, while in awake mice the %PF_awake-br_ was 80.3 ± 2.7 % (**Fig. 4B**).

**Figure 4.**
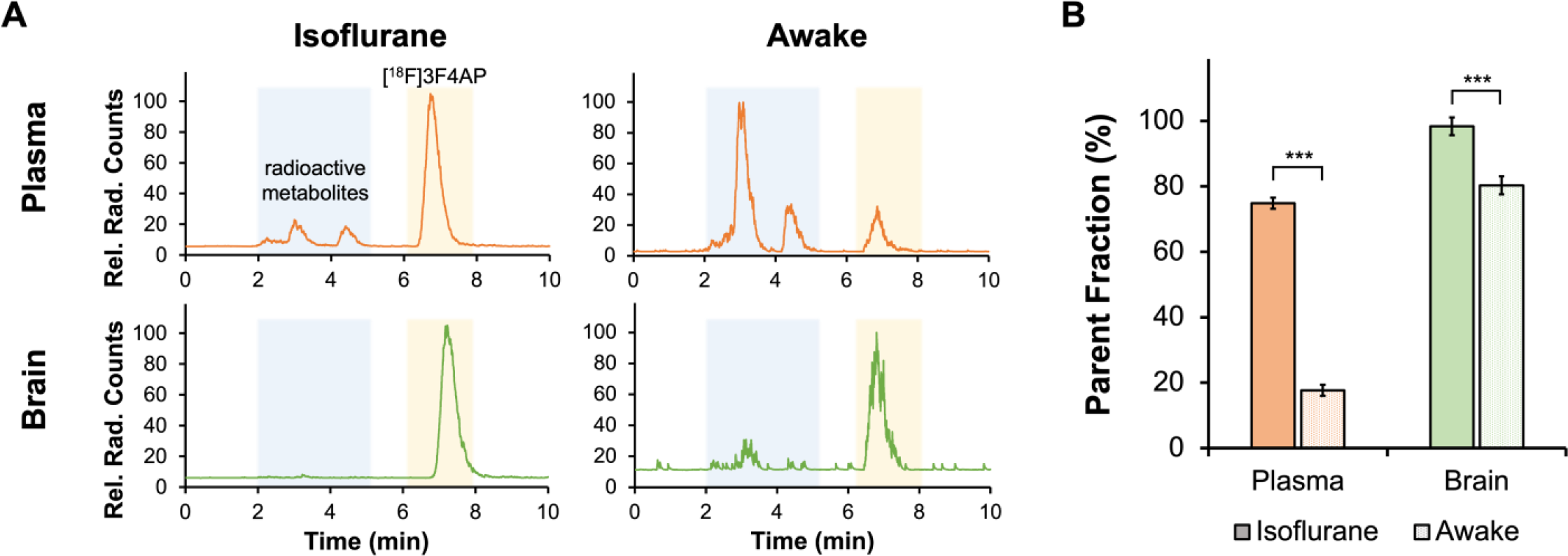
Radio-HPLC analysis of plasma and brain of anesthetized and awake mice. A) Representative radio-HPLC traces of plasma (orange) and brain (green) samples of anesthetized and awake mice. [^18^F]3F4AP parent peak signal is shown in yellow shaded box. Radioactive metabolites signals are shown in blue shaded box. B) Quantification of remaining percent parent fraction in plasma (%PF_iso-pl_ 74.8 ± 1.6 %, n = 10 *vs*. %PF_awake-pl_ 17.7 ± 1.7 %, n = 11; *p* < .001) and brain (%PF_iso-br_ 98.4 ± 0.2 %, n = 5 *vs*. %PF_awake-br_ 80.3 ± 2.7 %, n = 8; *p* < .001) after 35 min of [^18^F]3F4AP administration.

### Disulfiram treatment can partially rescue metabolic stability of [^18^F]3F4AP

The observed lower *in vivo* metabolism of [^18^F]3F4AP under isoflurane and our recent findings that 3F4AP is a good substrate for CYP2E1-mediated oxidation *in vitro* suggests that isoflurane and [^18^F]3F4AP can act as competing substrates for CYP2E1 *in vivo*. We postulated that in anesthetized conditions, isoflurane saturates CYP2E1 activity preventing metabolic breakdown of [^18^F]3F4AP by the P450 enzyme and resulting in an inflated metabolic stability of the radiotracer. Therefore, we sought to investigate if pharmacological inhibition of CYP2E1 could serve as a potential strategy to rescue [^18^F]3F4AP’s apparent metabolic stability observed under isoflurane anesthesia in awake subjects.

Disulfiram (DSF; Antabuse) has been used in the treatment of alcoholism for more than 60 years due to its ability to inhibit acetaldehyde dehydrogenase, which causes the build-up of acetaldehyde after ethanol consumption and causes an aversive reaction^27^. Resulting from its capability to inhibit several pharmacologically relevant target enzymes, a range of other therapeutic uses for DSF have been identified. CYP2E1 is one of the enzyme targets that can be inhibited by DSF. It has been shown that DSF administration inhibits CYP2E1 activity via mechanism-based inactivation of the P450 by an oxidized product of DSF formed *in vivo*^28^. DSF has been widely used as an inhibitor of CYP2E1 in various *in vivo* studies. For example, single-dose DSF has been proven to be an effective *in vivo* inhibitor of 2E1 in the hydroxylation of chlorzoxazone – a specific noninvasive probe of hepatic 2E1 activity^29^, and in the metabolism of volatile anesthetics such as enflurane^30^, sevoflurane^31^ and halothane^32^ in humans. Clinical isoflurane metabolism in humans has also been shown to be prevented by 80-90% with DSF treatment^33^. Preclinical work in rodents using CYP2E1 inhibition to reduce metabolic breakdown of pharmacologically active molecules has also been studied. For example, CYP2E1-mediated defluorination of the serotonin 1A receptor PET tracer [^18^F]FCWAY has been mitigated in rats with the use of miconazole – an antifungal agent and potent CYP2E1 inhibitor^34^. Inhibition of CYP2E1 to prevent defluorination of [^18^F]FCWAY was later extended in human imaging studies using DSF treatment, as a less toxic alternative to miconazole^35^.

To test the effect of DSF we selected a dose of 40 mg/kg, equivalent to 195 mg for a 60 kg human based on body surface area, which is within the clinical dose range for DSF (125 – 500 mg *q*.*d*.). A single dose of DSF 2h prior to radiotracer administration showed a 2.2-fold increase in the brain to blood concentration when compared to untreated awake mice (**Fig. 5A**). Additionally, presence of radioactive metabolites in the plasma of DSF-treated mice was reduced when compared to their untreated counterparts, evidenced by 50.0 ± 6.9 % of the parent [^18^F]3F4AP remaining in circulation at 35 min after tracer administration. Indication of metabolic breakdown in brain was nearly abolished with CYP2E1 inhibition, corroborated by similar unmetabolized [^18^F]3F4AP content in anesthetized and DSF-treated mice (%PF_iso-br_ 98.4 ± 0.2 % *vs*. %PF_DSF-br_ 95.0 ± 0.5 %) (**Fig. 5C**). Additionally, CYP2E1 inhibition was confirmed to be mediated by DSF treatment, with no significant differences observed between awake animals with and without 2h pre-treatment with sesame oil vehicle.

**Figure 5.**
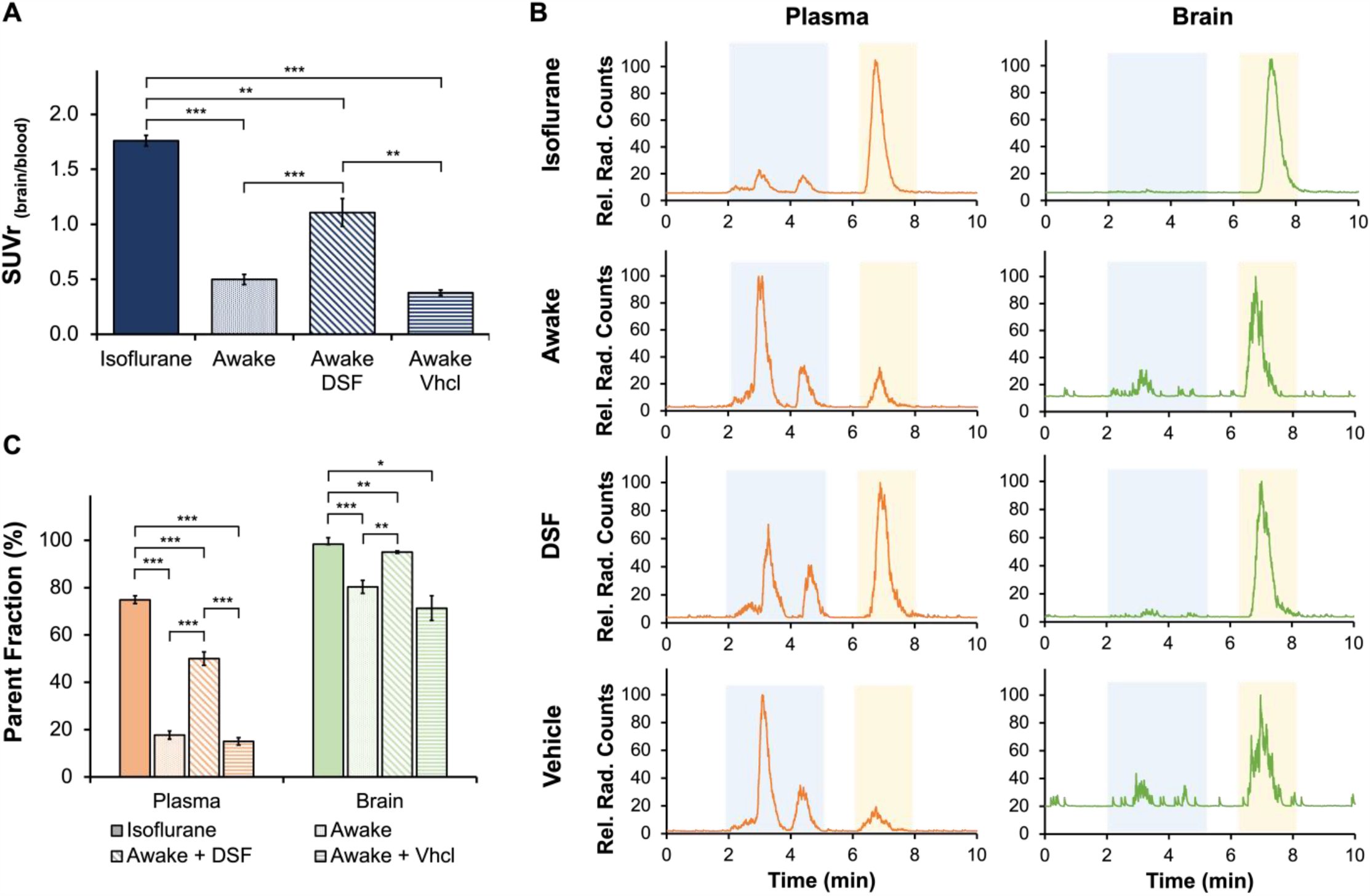
Inhibition of [^18^F]3F4AP metabolism *in vivo* with disulfiram treatment. A) Brain-blood ratio normalization of tracer uptake in whole blood and brain of anesthetized, awake, disulfiram-treated and vehicle control mice. B) Representative radio-HPLC traces of plasma (orange) and brain (green) samples of anesthetized, awake, and disulfiram-treated and vehicle control mice. [^18^F]3F4AP parent peak signal is shown in yellow shaded box. Radioactive metabolites signals are shown in blue shaded box. C) Quantification of remaining percent parent fraction in anesthetized, awake, disulfiram-treated and vehicle control mice in plasma and brain after 35 min of [^18^F]3F4AP administration.

## DISCUSSION

Our study demonstrates an artificially beneficial impact of isoflurane, the most commonly used anesthetic agent in animal PET studies, on the metabolism and brain uptake of the demyelination tracer [^18^F]3F4AP. Notably, the concentration of radioactive species in the brains of anesthetized animals was approximately 3.3 times higher than that in awake subjects. This effect is unexpected based on previous studies with [^18^F]FDG and a few other tracers that reported higher brain uptake in awake *vs*. anesthetized animals^36^. Furthermore, the high brain uptake in anesthetized monkeys and rodents had caused the expectation of higher brain uptake in humans than what was later found^9-11^. This unforeseen result prompts important questions about the underlying mechanisms responsible for this phenomenon.

A possible explanation for the increased brain uptake in anesthetized subjects is an increase in cerebral blood flow (CBF) under isoflurane. Even though it has been reported that isoflurane anesthesia can lead to an increase in CBF when compared to awake conditions^37^, the effects of isoflurane on CBF are dose-dependent and low concentrations of isoflurane like the concentrations used in this study tend to maintain CBF close to awake levels^38^. Therefore, it is more likely that an elevated bioavailability of the unmetabolized [^18^F]3F4AP in plasma, due to interference of isoflurane in the CYP2E1-mediated breakdown of the tracer, resulted in a greater transfer across the blood-brain barrier (BBB).

Our investigation into the metabolic stability of [^18^F]3F4AP yielded interesting outcomes. Although there was no significant difference in total radioactivity in the blood between anesthetized and awake mice, we detected substantial disparities in radiometabolite content in plasma. The higher concentration of metabolites in the plasma of awake compared to anesthetized mice (82.3% *vs*. 25.2%) strongly suggests that isoflurane has an inhibitory effect on the metabolic enzymes involved in [^18^F]3F4AP breakdown. Our research points towards CYP2E1 as a potential candidate enzyme, which has been previously reported to metabolize isoflurane and 4AP *in vivo* and has exhibited a high affinity for 3F4AP *in vitro*. Our results suggest that metabolites are not cleared from circulation more rapidly than the parent compound, or alternatively, the 35-minute time frame may not be sufficient to produce and eliminate metabolites.

We noted a substantially lower proportion of radiometabolites in the brain compared to blood, both in awake (18.3% *vs*. 82.3%) and anesthetized (1.6% *vs*. 25.2%) mice. This discrepancy suggests that metabolites do not enter the brain to the same extent as the parent compound. These findings align with a prior *in vitro* study that has characterized 5-hydroxy-3F4AP and 3F4AP N-oxide, the major and minor products of CYP2E1-mediated oxidation of 3F4AP, to exhibit considerably lower permeability across an artificial brain membrane compared to 3F4AP. These two oxidation products are likely the *in vivo* metabolites of [^18^F]3F4AP, which permeate the brain at a much lower rate than the unmetabolized compound. Another possibility is the production of these metabolites within the brain, though CYP2E1 expression in the brain seems to be negligible in basal conditions^39^. Although we cannot entirely exclude the possibility of some metabolites crossing the BBB, a large fraction of the detected brain metabolites is likely a result of metabolites present in remaining blood in the vascular space of the brain, since perfusion was not performed, and minimal metabolism occurs within the brain.

Our study presents a promising avenue for mitigating the *in vivo* metabolic breakdown of [^18^F]3F4AP. We observed that a single dose of disulfiram, a known CYP2E1 inhibitor, could partially rescue the metabolic stability of the tracer. Importantly, the dose of disulfiram we used was equivalent to that employed in humans. Given the potential utility of disulfiram in limiting metabolism in human imaging, it holds promise as a strategy to enhance the reliability and consistency of [^18^F]3F4AP studies in future imaging investigations.

The impact of our study likely goes beyond [^18^F]3F4AP. Most PET tracers are metabolized to some degree and CYP2E1 is the most abundant P450 enzyme in the liver and likely responsible for metabolizing other small molecule tracers. Even though isoflurane has been reported to be metabolized by CYP2E1, the ability of isoflurane to saturate the enzyme thus preventing tracer metabolism has not been previously recognized. Instead, differences encountered between animal and human studies have commonly been vaguely ascribed to species differences^12^. Auspiciously, this work describes a simple experimental method that can be used to test the effect of anesthesia on tracer metabolism and brain uptake before translating tracers to humans.

## ACKNOWLEDGEMENTS

We thank David Lee and Kyle Stewart at the MGH Gordon PET cyclotron facility for producing fluorine-18. Funding: This study was supported by R01NS114066 (PB), S10OD018035, K99EB033407 (YPZ) and MGH-ECOR PSDA award (KMRT).

## AUTHOR CONTRIBUTIONS

KMRT contributed to the study design, performed the radiosynthesis of the tracer, conducted the animal studies and analyzed the data; YS performed the blood processing and radiometabolite analysis; KT and YPZ: contributed in the animal studies. PB contributed to the study design, data analysis and interpretation. KMRT and PB wrote the manuscript and all authors reviewed and approved it.

## DATA AVAILABILITY

The datasets generated and/or analyzed during the current study are available from the corresponding author upon reasonable request.

